# Substantial genomic and methylation variability between MCF-7 sublines

**DOI:** 10.64898/2026.02.17.706500

**Authors:** Halimat C Atanda, Adam D Ewing

**Affiliations:** Mater Research Institute - University of Queensland

## Abstract

Cancer cell lines have long been used as *in vitro* models for molecular assays in diagnostic and therapeutic development due to their accessibility as a well-controlled system. MCF-7 cell lines are the most widely studied cell lines in human breast cancer research, and its sublines have been reported to exhibit clonal, cytogenetic, and transcriptomic variability. However, allele-specific methylation alterations in cancer genomes remain inadequately explored, largely due to limitations in sequencing methods. Here, we applied nanopore sequencing technology to characterise the genomic and epigenomic landscapes of two MCF-7 sublines. We identified global and local DNA methylation differences as well as structural variants (SVs), and single-nucleotide variants (SNVs) between and within the sublines. Our analysis revealed substantial divergence in methylation patterns between the sublines, with ∼3% of the differentially methylated regions (DMRs) overlapping with known cancer driver genes. These DMRs overlap breast cancer-associated genes, including ERBB2, CDH1, SALL4, GATA2, GATA3, HMGA2, and FBLN2. We find that the majority of differentially methylated sites are explained by differential allelic methylation, and that allele-specific DMRs often overlap points where antisense non-coding RNAs overlap protein-coding genes. Transposable elements in both sublines also showed distinct methylation profiles, with one subline having hypomethylated L1 elements compared to the other, which correlated with the amount of apparent insertional mutagenesis attributable to L1 between the sublines. Our study demonstrates the utility of nanopore sequencing in providing novel insights into genomic and methylomic differences within cell lines, in addition to insight into the nature of differential allelic methylation.

## Introduction

Human cell lines have long served as an accessible, controlled environment to study molecular interactions in cancer. The MCF-7 cell line, established in 1973 at the Michigan Cancer Foundation from the pleural effusion of a 69-year-old woman with metastatic breast carcinoma, is the most widely studied model in human breast cancer research (Comsa et al., 2015; Soule et al., 1973). This cell line has contributed to understanding breast cancer pathology, drug development, and mechanisms of therapeutic resistance (Comsa et al., 2015). However, the wide distribution and diverse culture conditions of MCF-7 cells across research groups and laboratories have led to the emergence of sublines with different proliferation and differentiation rates, affecting research reproducibility and clinical translation (Kleensang et al., 2016; Nugoli et al., 2003; Osborne et al., 1987). For instance, Osborne et al. (1987) compared four MCF-7 sublines and reported structural chromosome alterations as well as differences in hormone content and growth rate. Nugoli et al. (2003) also reported divergence in eight MCF-7 sublines at the genomic and transcriptomic level, with the ATCC subline sample being highly divergent and having fewer losses and gains compared to the other 7 sublines. These differences, also reported in drug response studies (Ben-David et al., 2018), underscore the importance of understanding subline-specific genomic and methylation alterations that could influence experimental outcomes.

DNA methylation is the closest epigenetic layer to the DNA sequence (Liang & Weisenberger, 2017; Nebbioso et al., 2018). Many methods for its ascertainment have been devised, often involving conversion of unmethylated cytosine to uracil via chemical or enzymatic means. This has disadvantages when it comes to repetitive regions of the genome and when attempting to assess allele-specific methylation. Nanopore sequencing generates long to ultra-long read sequences that can exceed 1Mbp, and simultaneously detects chemical modifications to sequenced bases, including DNA methylation. This ability offers an opportunity to comprehensively investigate genetic variants and methylation differences in cancer genomes (van Dijk et al., 2018; Weirather et al., 2017) while overcoming the limitations of previous methods like whole genome bisulfite sequencing and microarrays, which can include DNA damage and restricted coverage of the genome.

In this study, we use nanopore sequencing technology to explore genomic and methylomic differences between MCF-7 cell lines originating from two geographically separated cell line distributors: the American Type Culture Collection (ATCC) and European Collection of Authenticated Cell Culture (ECACC). This study aims to provide an understanding of subline-specific epigenetic landscapes in this widely used breast cancer model that may be essential for translational studies.

## Methods

### Sequence data processing

Oxford Nanopore Technologies sequence data from MCF-7 cells was obtained from NCBI SRA accession PRJNA748257 (Cheetham et al 2022). Raw current signal data in fast5 format were converted to nucleotide bases using guppy basecaller version 6.2.1 (and minimap2 version 2.22-r1101) (Wick et al., 2019) with model configuration file, dna_r9.4.1_450bps_modbases_5mc_cg_sup.cfg for PromethION. The basecalled reads were then mapped to the human reference genome GRCh38 with the guppy –align option and saved in bam output with modified base tags. The aligned reads for each subline were sorted using samtools version 1.15 and saved in separate bam files for downstream analysis.

### Identifying and visualising DNA methylation

We used the Bioconductor R package, Dispersion Shrinkage for Sequencing data (DSS version 3.17) for differential methylation analysis in both sublines (Wu et al., 2015). DSS uses the beta-binomial distribution to model the methylation level at each CpG site, based on methylation counts, and then uses a Wald test statistic on the coefficients of the distribution to identify regions with differential methylation (Feng & Wu, 2019). DSS expects an input file containing chromosome number, genomic coordinates, number of reads, and number of methylations, for which we used methylartist “wgmeth” to generate from the bam files of each subline. We used the DMLtest function with the smoothing option set to true to estimate the mean loci methylation between the sublines, and the callDMR function to identify differentially methylated CpG regions (DMRs).

Candidate DMRs were ranked by areaStat, which aggregates significant DML calls into regions. Selected DMRs were manually examined with the integrative genome viewer (IGV) and visualised with methylartist version 1.2.6 (Cheetham et al., 2022). Using methylartist “locus”, we generated images for selected DMRs. Methylartist “segmeth” and “segplot” were used to compare the global methylation profile of the ATCC and ECACC sublines by segmenting the genome into 10kbp bins via bedtools (Quinlan and Hall 2010). Additionally, methylartist segmeth and segplot were used to generate methylation plots for transposable element (TE) subfamilies in both sublines: ALU, SVA, and L1. Methylartist “region” was used to compare the methylation profiles across the entire chromosomes of both sublines.

### Variant/mutation calling

Sniffles version 2.0.7 (Sedlazeck et al., 2018) was used to identify structural variants between the sublines and classify them into insertions, deletions, and other forms of chromosomal rearrangements. We filtered the resulting variant file to retain only the variant calls that passed all quality control filters and were specific to either cell line.

We used tldr version 1.2.2 (Ewing et al., 2020) to detect and annotate non-reference transposable elements (TEs) in long read alignments. Insertion calls from tldr were considered if they were annotated as passing all filters, were specific to one subline, and novel as indicated by “NA” in the NonRef column. Insertions were further filtered by manual examination with BLAT on the UCSC Genome Browser (Kent et al 2002).

We performed somatic single nucleotide variant (SNV) calls with ClairS (Zhenxian et al., 2023) to detect somatic small variants. Variants were called twice, such that each subline was analysed as both the tumour and normal sample. The variant files were filtered to retain passed calls and annotated with gnomAD allele frequencies to eliminate known germline SNVs. Variant calls were filtered to retain only somatic and subline-specific variants.

### DMR Annotation

DMRs were annotated with the subline-specific mutation calls generated here, and with human genome (hg38) regulatory annotations including those from GeneHancer (Fishilevich et al 2017) and ENCODE cis-regulatory elements (CREs) (ENCODE Project Consortium et al. 2020), which include CTCF sites, DNase hypersensitive sites, H3K4me3 (K4m3) sites, distal enhancer-like signatures (enhD), proximal enhancer-like signatures (enhP), and promoter-like signatures (prom). We used the Cancer Gene Census, downloaded from the COSMIC database (Sondka et al 2018) in April 2023, and significantly mutated genes in the TCGA Breast Cancer dataset, downloaded from the Xena Browser (Goldman et al 2020) database in December 2024, to identify known cancer genes overlapping each DMR. We created custom Python scripts for the annotation and used cyvcf2 (Pedersen and Quinlan, 2017) to parse variant call files (VCFs). All VCFs were annotated using Ensembl Variant Effect Predictor (VEP v111) (McLaren et al., 2016).

### Haplotagging

MCF-7 variants previously derived from Illumina sequencing were phased with long read data using Whatshap version 0.18 (Martin et al., 2016). We used the whatshap “haplotag” and the phased germline variants to tag alignment reads from each subline by haplotype, resulting in a haplotagged bam file for each subline. The methylartist command wgmeth was run with the --phased and –dss options to generate DSS-compatible input files. Haplotype-specific DMRs and DMLs within each subline were called via DSS, as described above. Haplotype-specific DMR annotation was carried out as described for the unphased sequence data.

## Results

### Widespread differential methylation between sublines

Examination of overall CpG methylation indicated that one subline (ATCC derived) is hypomethylated in comparison to the other subline (ECACC derived) (Figure 1, Figure 2a). Visualisation of methylation levels across chromosomes (Figure 1) shows differential methylation, with larger divergences localised to distinct regions of chromosomes and smaller differences spread across the entire chromosome. In considering ENCODE cis regulatory elements (CREs), we observed higher methylation of CREs in the ECACC subline (Figure 2b), particularly at proximal and distal enhancers (enhP and enhD, respectively), at CTCF sites, and H3K4me3 sites.

**Figure 1.**
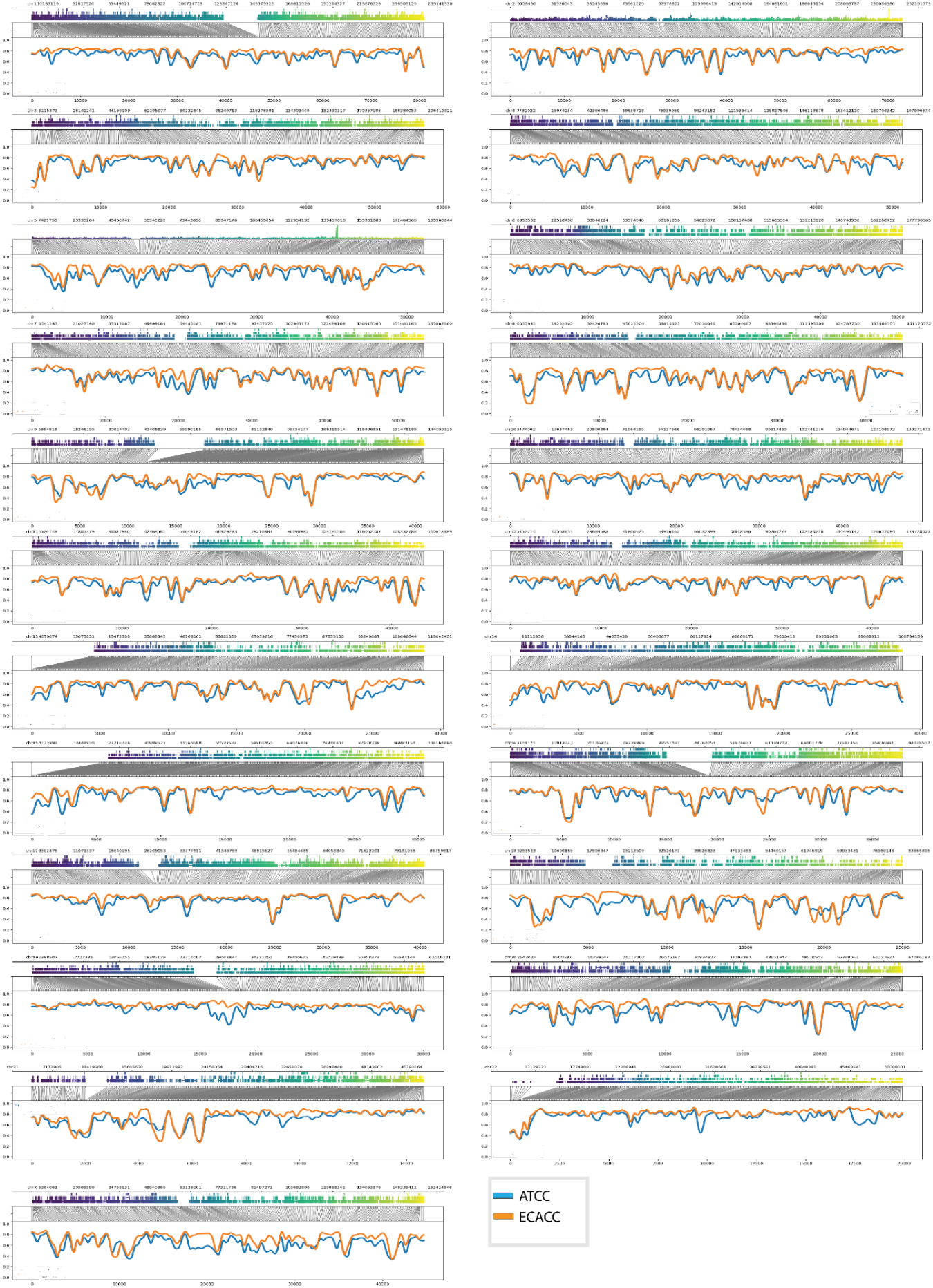
Methylation profiles for whole chromosomes in MCF-7 sublines in comparison. Each panel shows genome coordinates, distribution of exons and introns, translation from genome coordinates to modified bases, and smoothed plot of methylation profile coloured by sample, in that order.

**Figure 2.**
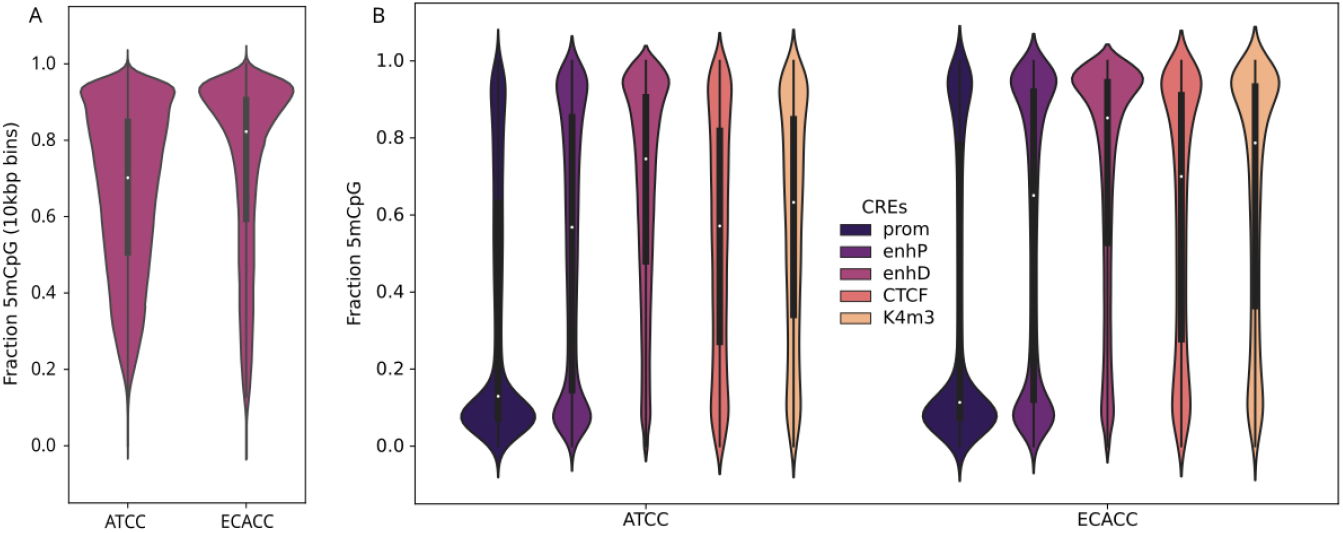
Violin plots of cis-regulatory elements CpG methylation **(A)** and aggregate whole genome CpG methylation of both MCF-7 sublines in 10kbp intervals **(B)**.The y-axis represents the methylation percentage and shows the bulk of CpG sites in the ECACC subline are hypermethylated compared to the ATCC subline which has CpG sites with a wide range of methylation levels. Similarly, all regulatory elements in the ECACC sublines are hypermethylated compared to the ATCC subline.

We next sought to examine differential methylation at specific sites via statistical analysis using DSS, which yields differentially methylated loci (DMLs) and their aggregation into differentially methylated regions (DMRs). We ranked DMRs by area statistics (areaStat), which aggregates the test statistics of individual CpGs. Since areaStat values have no standard cutoff value, we defined DMRs as those with absolute “areaStat” values above 1000.

In total, excluding unplaced contigs and alternate haplotypes, we identified 1909 DMRs between the two sublines (Supplemental Table 1, areaStat > 1000), of which 56.5% (n = 1079) were in ENCODE-annotated promoters. We note that as both samples have relatively similar coverage (x28 ATCC vs x30 ECACC), these differences are unlikely to arise from differences in sequencing coverage, which may influence methylation call accuracy (Liu et al., 2021).

From the phased data (Supplemental Table 2), the methylation distribution between the haplotypes of each subline showed the ECACC subline had substantially fewer allele-specific DMRs (435, areaStat > 1000, Supplemental Table 2a) compared to the ATCC subline (959, areaStat > 1000, Supplemental Table 2b), indicating the ATCC subline was more prone to haplotype-specific methylation alterations. In both sublines, allele-specific DMRs were widely distributed across most chromosomes, except for chromosomes 18 and 21, which had relatively lower numbers of allele-specific DMRs in both sublines, possibly due to aneuploidy. Allele-specific DMRs corresponded to ENCODE-annotated promoters in 285 (65.5%) and 622 (64.8%) of DMRs for ECACC and ATCC sublines, respectively.

In general, we find that the majority of DMRs between cell lines are due to differences that affect a single allele. As a sample, we selected a subset of 98 DMRs (those that intersect genes in the Cancer Gene Census), which, after curation and merging of overlapping windows, yields a set of 63 DMRs (Supplemental Table 3). Of those 63, 13 appear to be driven by differences on both alleles, and 31 are driven by differences in the CpG methylation of one allele, with the remaining DMRs being ambiguous due to insufficient phased sequence coverage.

### Allele-specific differential methylation of breast cancer-associated genes

At the time of retrieval, the cancer gene census data contained 737 genes, of which 46 were associated with breast cancer (BC). DMRs identified between the ATCC and ECACC sublines overlapped at least 7 BC associated genes: GATA3, ERBB2, FBLN2, GATA2, SALL4, CDH1, and HMGA2 (Fig 3, Fig S1-2, Fig S3a-6a). All of these genes are either known oncogenes or tumour suppressor genes, or function in both capacities. All DMRs overlapping BC genes contained one or more ENCODE candidate cis-regulatory elements, including proximal and distal enhancer-like signatures and promoter-like signatures.

**Figure 3.**
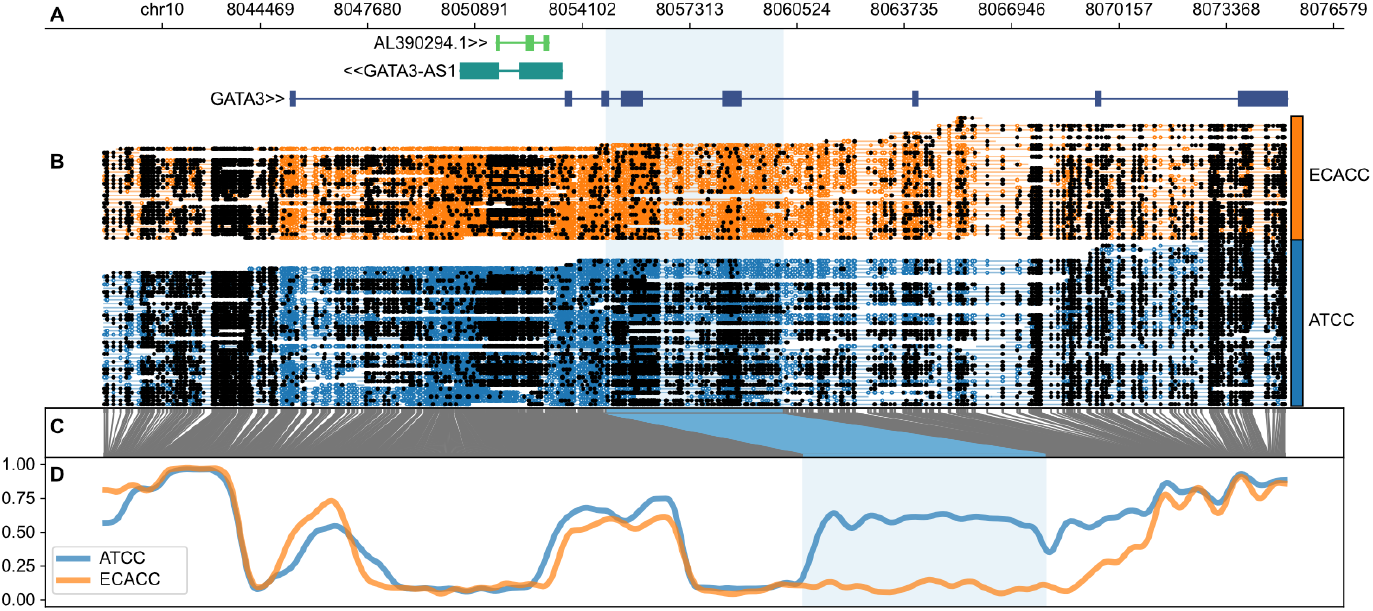
GATA3 as a representative example of a DMR overlapping a gene. Panel **(A)** shows genome coordinates (hg38) with Ensembl gene annotations depicted as thick boxes depicting the mRNA with thin lines depicting introns. The direction of transcription is annotated next to the gene name. Panel **(B)** shows read alignments with coloured dots representing unmethylated CpGs and filled dots representing 5mCpGs. Correspondence between read alignment blocks and samples is shown to the right of the alignments. Panel **(C)** shows the translation from genome space into a CpG-only space, which permits efficient plotting of the methylation pattern shown in panel **(D)** at the bottom. The highlighted region indicates the DMR of interest in the plot.

GATA3 is a transcription factor critical for maintaining luminal cell identity in breast tissue that is often altered in breast cancer. Studies have linked high or differential over-expression of wild-type GATA3 in ER+ breast cancers with inhibited metastasis, high chances of disease-free survival, and overall better prognosis (Afzaljavan et al., 2021; Takaku et al., 2015; Yan et al., 2010). GATA3 is frequently mutated in breast cancers, including TCGA samples (Cancer Genome Atlas Network 2012), and some of its mutations are associated with tumorigenesis, estrogen receptor (ER) signalling, and metastasis. It has also been speculated that GATA3 wild-type expression may contribute to maintaining a phenotype with favourable prognosis (Afzaljavan et al., 2021). We find that the differential methylation pattern of GATA3 (Fig 3f) is driven by allele-specific hypermethylation on one allele within the ATCC-derived line (Fig 4).

**Figure 4.**
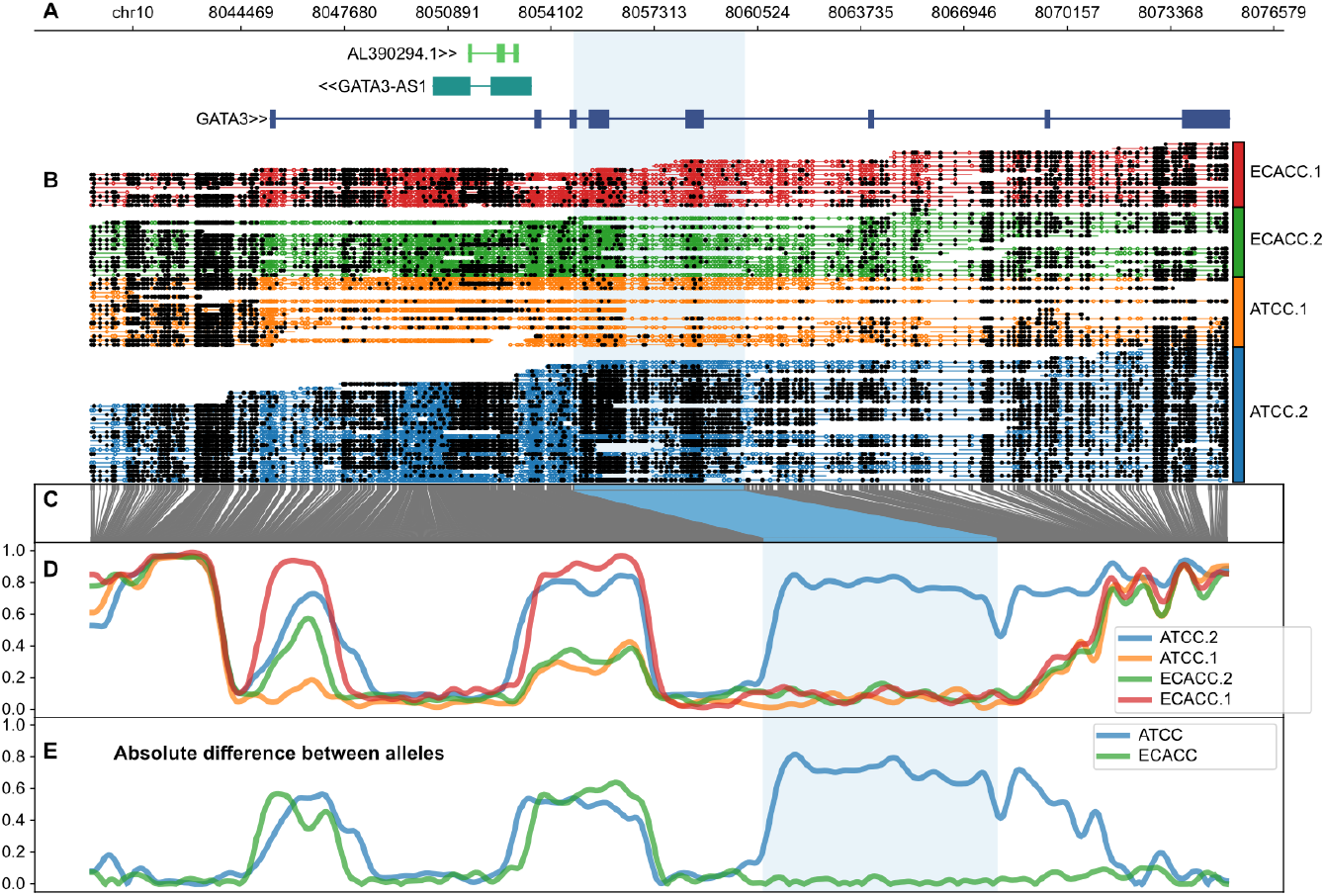
Allele-specific methylation of the GATA3 region shown in Figure 3. Panels A-D have the same interpretation as per Figure 3, with samples split into haplotypes 1 and 2 as shown as a suffix on the sample names. The additional panel **(E)** at the bottom shows the absolute difference between the two alleles for each sample as indicated.

We see similar patterns where an overall differential methylation pattern between lines is driven by allele-specific methylation in one line in additional cancer-specific genes. These include GATA2 (Fig S3a-b) which is linked to poor prognosis and may downregulate the tumour suppressor gene PTEN in breast cancer (Wang et al 2012), SALL4 where overexpression is oncogenic in breast cancer (Reviewed in Dirican and Akkiprik 2016) (Figs S4a-b), CDH1 where mutations are a major risk factor for lobular breast cancer (Pharoah et al 2001) (Fig S5a-b), and HMGA2 (Fig S6a-b) which is highly expressed in malignant breast tumours (Sun et al 2013) and is associated with poor outcomes (Wu et al 2016).

In general, we find that differential methylation is very often driven by hyper- or hypomethylation of one allele. Additional examples of genes implicated as cancer drivers include MYCN (Fig S7a-b), which is overexpressed across many cancers, including triple-negative breast cancer (Schafer et al 2020), FGFR2 (Fig S8a-b), and COL2A1 (Fig S9a-b). A majority of DMRs overlapping with genes in the Cancer Gene Census (CGC) are differentially methylated in an allele-specific manner (Supplemental Table 3), highlighting the importance of long read sequencing technologies, which enable allelic resolution combined with 5mC resolution.

Examining the allele-specific methylation pattern of DMRs overlapping with genes in the CGC, we find that many overlap with antisense RNAs: GATA3/GATA3-AS1 (Figs 3-4) MYCN/MYCNOS (Fig S7a-b), LEF1/LEF1-AS, EBF1/LINC02202, CDK6/CDK6-AS1, ZEB1/ZEB1-AS1, WT1/WT1-AS, HOXC11/HOTAIR, NKX2-1/NKX2-1-AS1 (Figs S10-S16). These instances seem similar to the IGR2R/AIRN locus, where transcription of AIRN induces imprinting (parent-of-origin specific expression) of IGF2R (Latos et al 2012, Santoro et al 2013), and is perhaps consistent with co-clustering of ncRNAs with allele-specific expression (ASE) of protein-encoding mRNAs (Hasenbein et al 2025).

### Differential mutation burden

We identified 66 candidate subline-specific TE insertions, 5146 candidate subline-specific SVs, and 26003 candidate subline-specific SNVs. Removing variants on unplaced contigs yielded 4774 SVs and 60 TE insertions. Removing calls likely due to SVA VNTR expansions (Ewing et al. 2020) yielded the 53 insertions shown in Supplemental Table 4 followed by further manual examination of TE insertions using BLAT (UCSC Genome Browser) identified 25 “high confidence” insertions (HC TEs) (see “High Confidence” column in Supplemental Table 4). The ATCC subline sample consistently had a higher number of subline-specific variants (4348 SVs, 16777 SNVs, and 24 HC TEs) compared to the ECACC subline (426 SVs, 9226 SNVs, and 1 HC TE).

Using VEP with default settings, 7606 (45%) and 6550 (71%) of subline-specific SNVs were found to be novel in ATCC and ECACC sublines, respectively, and both sublines largely contained intronic variants. The distribution of coding region variants was different between sublines, with the ATCC subline having more missense variants and fewer stop gains (69% and 1%, respectively) compared to the ECACC subline (55% and 6%, respectively).

Both sublines had a similar distribution of structural variant types, with insertions accounting for at least 50% and deletions, at least 40%. More than two-thirds (70%) of the ATCC subline variants involved coding sequences, and 38% were intronic structural variants. In the ECACC subline genome, 94% of the structural variants potentially impacted coding sequences, and 79% represented transcription factor binding site (TFBS) variants, amplification and ablation. The prominence of structural variants in the TFBS of the ECACC subline suggests that its transcriptional regulation may be substantially altered. Considering the critical role of TFBSs in regulating gene expression (Iñiguez-Muñoz et al., 2024; Moore et al., 2013), disruption by structural variants could affect TF binding and subsequently lead to altered gene expression.

### Differential methylation of TEs and differential insertional mutagenesis

In somatic cells, methylation is the primary means by which retrotransposition activity is controlled (Greenberg and Bourc’his 2019). We analysed methylation distribution across active TE families Alu, SVA, L1, and the L1 5’ UTR, which controls L1 expression genome (Fig 5). We found higher level of L1 methylation across multiple subfamilies including the active L1Hs subfamily (Fig S17) in the ECACC subline compared with lower 5mCG on L1 elements in the ATCC genome, which is generally consistent with the observation of higher L1 element activity in the ATCC genome (Supplemental Table 3), as well as Alu activity which is driven *in trans* by L1 (Dewannieux et al 2003). Both sublines showed high methylation (fraction between 0.8 and 1.0) on Alu and SVA elements.

**Figure 5.**
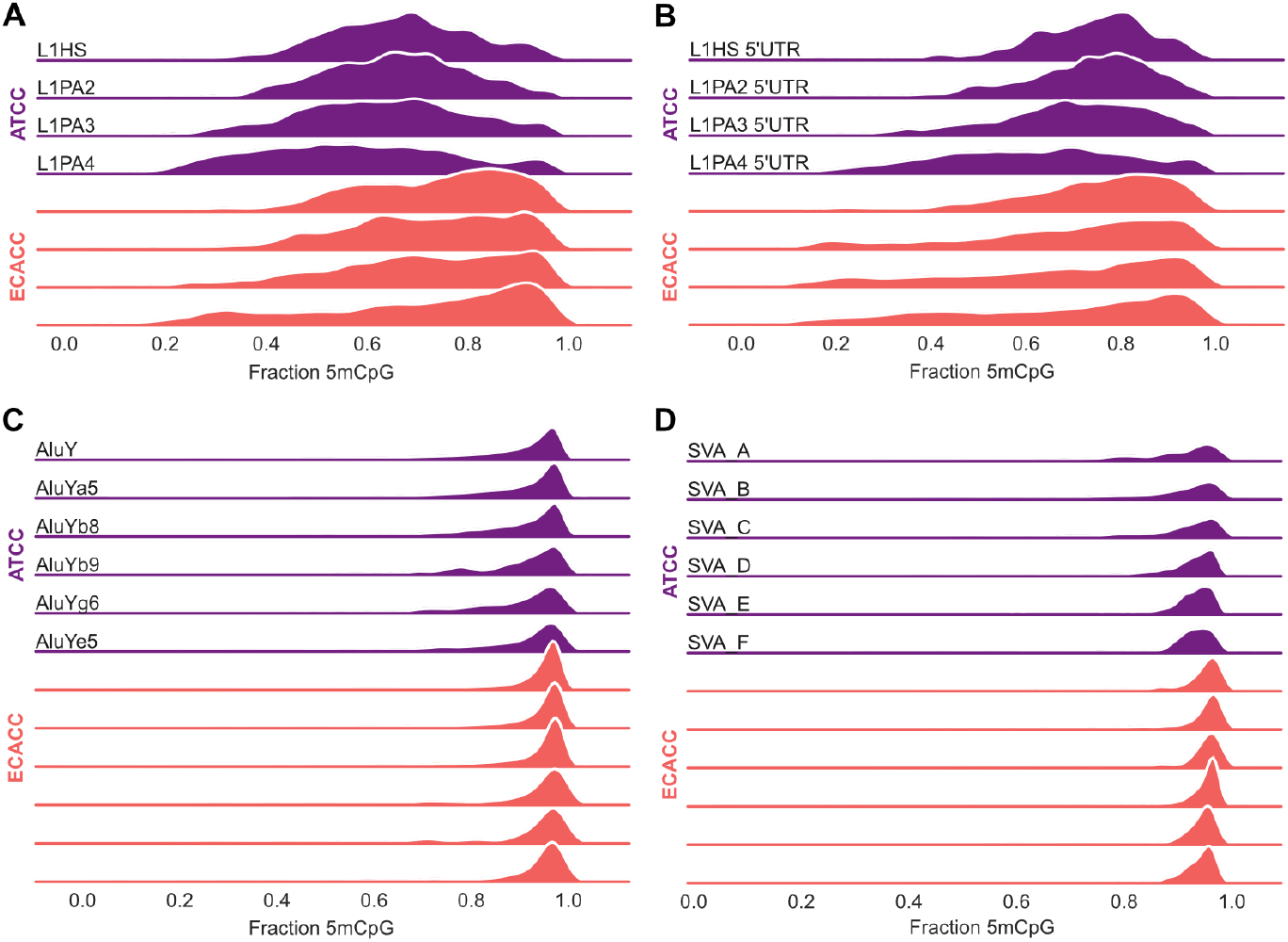
Ridge plots for CpG methylation in retrotransposons, including long interspersed element 1 LINE-1 families, Alu and SVA, compared between sublines. The ridges represent the distribution of methylation for each subfamily. The range on the x-axis corresponds to the level of methylation, where 0 represents no methylation and 1 represents complete methylation. Full-length L1 elements **(A)** and their CpG-rich 5’ UTRs **(B)** across multiple subfamilies show substantial differential methylation between ATCC and ECACC derived cell lines. This differential methylation is also present in Alu **(C)** and SVA elements **(D)** from various subfamilies to a lesser extent.

Cancer cells often exhibit hypomethylation at repetitive elements, especially the L1 regions, and this feature has been associated with poor clinical outcomes (Park et al., 2014; Xiao-Jie et al., 2016; Zeggar et al., 2020). The reduced L1 methylation in the globally hypomethylated ATCC subline is consistent with studies reporting L1 hypomethylation to be closely correlated with global hypomethylation (Barchitta et al., 2014; Pappalardo & Barra, 2021), which is often used as a marker of genome-wide methylation given its abundance (up to 17%) within the human genome. Philippe et al. (2016) reported high and locus-specific L1 expression and the presence of non-reference L1 insertions in MCF-7 cells coupled with their ability to permit selective escape from regulatory mechanisms, like epigenetic and post-transcriptional controls.

While Alu repeats are hypomethylated in several cancers, including colorectal, gastric, and breast cancer (Barchitta et al., 2014; Park et al., 2014), this does not appear to be the case in MCF7. Although our data show high Alu and SVA methylation in both subline samples, these repeat elements might still be hypomethylated when compared to non-cancer cells. Also, Alu and SVA repeats may still be transcriptionally active in the ATCC subline as they rely on L1 for propagation (Hancks et al 2011). Overall, the ATCC subline could have higher genomic instability as a consequence of its hypomethylated L1 elements contributing to the higher observed somatic retrotransposition load.

## Discussion

MCF-7 cell lines exhibit genomic instability associated with abnormal gene expression and varying phenotypes, including response to treatment (Ben-David et al., 2018; Nugoli et al., 2003). Epigenetic modifications, especially aberrant DNA methylation, are established pathogenic hallmarks in all cancers. In this study, we report genomic differences in terms of mutation burden and epigenomic differences in terms of DNA 5mC methylation alterations, obtained using nanopore sequencing technology. Our findings support previously reported high levels of genetic variability between MCF-7 sublines, potentially driven in part by global methylation differences as observed for TE insertion activity.

We identified differentially methylated breast cancer genes between and within the sublines, underscoring the potential impact of methylation alterations in regulating key pathways for oncogenes and tumour suppressors. The genes identified in each subline highlight the distinct epigenetic landscapes that may arise from subline-specific genomic aberrations or clonal evolutionary pressures. GATA3, whose expression has been reported to be positively correlated to BRCA1 expression to control metastasis (Afzaljavan et al., 2021; Bai et al., 2021), was differentially methylated between sublines and may serve as a relevant marker of genomic instability. Using nanopore sequencing data, we were able to explore and report differential allele-specific methylation patterns on GATA3, among many other genes, between the sublines included in this study.

Aberrant methylation is either an abnormal increase (hypermethylation) or decrease (hypomethylation) in 5mC content, and hypomethylation is generally expected to impact larger regions of the genome than hypermethylation (Pappalardo & Barra, 2021). Studies by Nugoli et al. (2003) and Osborne et al. (1987) reported the ATCC subline to be genomically unstable or divergent in comparison with other sublines, supporting our findings of high mutational burden, overall global hypomethylation, and hypomethylation around repeat elements, specifically L1s, within the ATCC subline. L1 elements are known contributors to genomic instability, and their reactivation in cancer cells seems to have implications for disease progression (Xiao-Jie et al 2016). The potentially increased L1 activity in the ATCC subline could disrupt gene expression neighboring insertion sites and L1-induced double-strand breaks can lead to deletions and chromosomal rearrangements (Kazazian 2004). The combined effect of these genomic and methylomic differences could result in a range of impacts, including dysregulated TF binding leading to altered gene expression, and possible aggressive tumour behaviour due to altered cellular protein production (Bhandari et al., 2023; Gebhard et al., 2010; Moore et al., 2013).

Compared to methylation within gene bodies, the role of methylation in promoter and regulatory regions is better understood, where hypermethylation reduces the expression of the gene and vice versa. Studies have suggested gene body methylation can maintain transcription accuracy, prevent TF binding, and regulate alternative splicing (Campagna et al., 2021; Wang et al., 2022). The functional implication is often dependent on the location of the aberrant methylation (intron or exon), the role and mechanism of action of the affected gene, as well as the tissue or cell in which the aberrance occurs. Therefore, while it may not be apparent how gene body differential methylation in GATA3 and ERBB2 may affect tumour behaviour in each subline in terms of invasiveness or therapy response, the differential methylation patterns observed between the sublines show the dynamic nature of the cancer epigenome and its likely contribution to phenotypic diversity. Future studies may focus on validating allele-specific methylation changes, keeping in mind that methylation patterns in cell lines will vary from methylation patterns in primary tissues or cells, and elucidating their functional consequences, potentially contributing to the development of epigenetic therapies.

## Acknowledgements

We acknowledge the Translational Research Institute (TRI) for research space, equipment, and facilities. This study was funded by the Australian Department of Health Medical Research Future Fund (MRFF) (MRF1175457 to A.D.E.), and by support from the Mater Foundation.

## Data Availability

Nanopore sequence data from MCF7 lines was obtained from NCBI SRA accession PRJNA748257. Variant calls are available at figshare (10.6084/m9.figshare.31360360).

## List of Abbreviations

5mC/5mCpG/5mCG: 5-methyl cytosine, 5-methyl cytosine-guanine
AS: Antisense
ASE: Antisense expression
ATCC: American Type Culture Collection
CDH1: Cadherin-1 / Epithelial Cadherin
CDK6: Cyclin-dependent kinase 6
COL2A1: Collagen type II alpha 1 chain
CpG: Cytosine followed by Guanine in a DNA context
CRE: cis-regulatory element
DMR: Differentially Methylated Region
EBF1: Early B-cell factor 1
ECACC: European Collection of Authenticated Cell Cultures
ERBB2: Erb-B2 Receptor Tyrosine Kinase 2, aka HER2
FBLN2: Fibulin-2
FGFR2: Fibroblast growth factor receptor 2
GATA2: GATA-binding protein 2
GATA3: GATA-binding protein 3
HC: High-confidence
HMGA2: High mobility group AT-hook 2
HOTAIR: HOX Antisense Intergenic RNA
HOXC11: Homeobox C11
L1 / LINE-1: Long INterspersed Element-1
LEF1: Lymphoid enhancer-binding factor 1
mRNA: messenger RNA
MYCN: N-myc / basic helix-loop-helix protein 37
ncRNA: non-coding RNA
NKX2-1: NK2 homeobox 1
PTEN: Phosphatase and tensin homolog
SALL4: Spalt-like transcription factor 4
SNV: Single-nucleotide variant
SV: Structural variant
TE: Transposable element
TF: Transcription factor
TFBS: Transcription factor binding site
VCF: Variant call format
VNTR: Variable number of tandem repeats
WT1: Wilms’ tumour-1
ZEB1: Zinc finger E-box binding homeobox 1

