## Supplementary Figures S1-S17 for "Substantial genomic and methylation variability between MCF-7 sublines"

### Supplemental Figures S1-S17

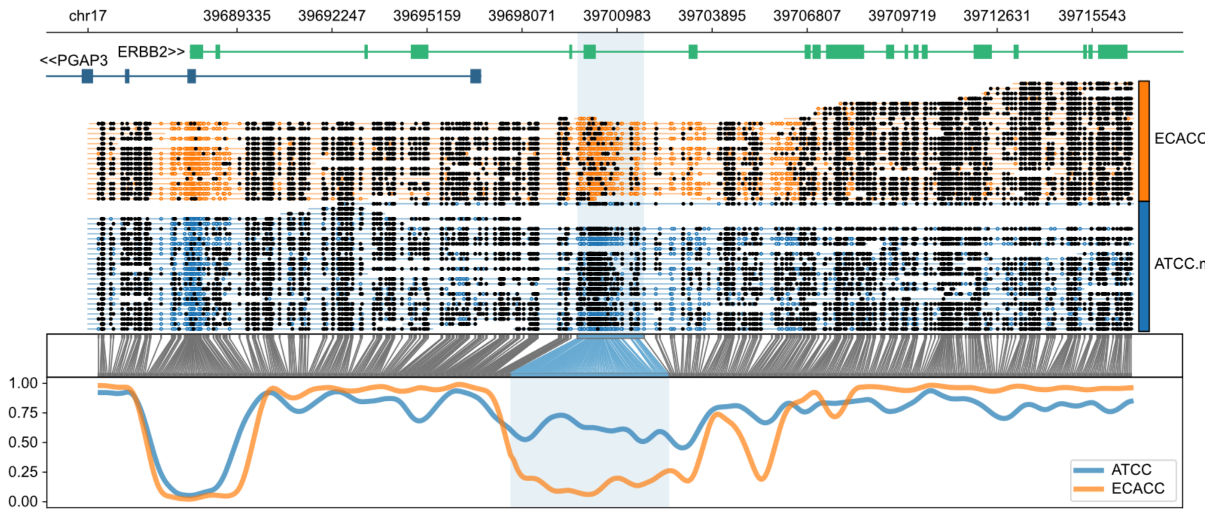


**Supplementary Figure 1:** ERBB2 differential methylation pattern

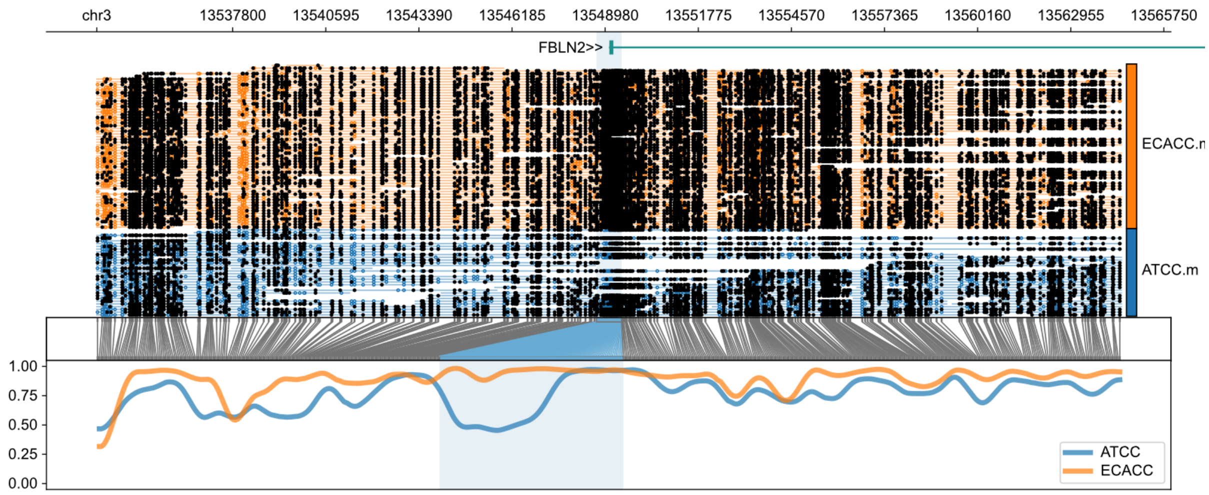


**Supplementary Figure 2:** FBLN2 differential methylation pattern

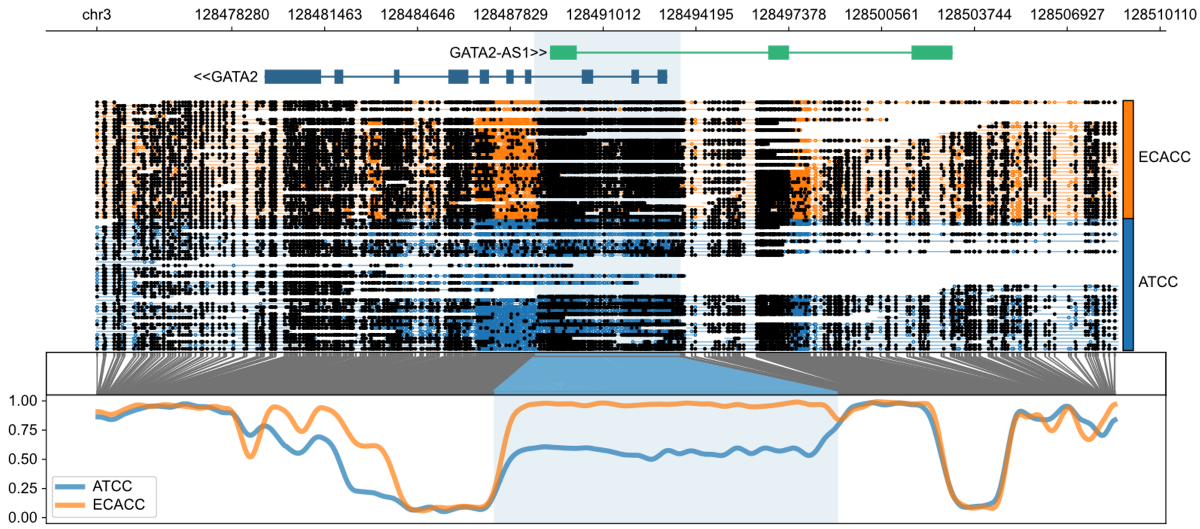


**Supplementary Figure 3a:** GATA2 differential methylation pattern, not allele-resolved

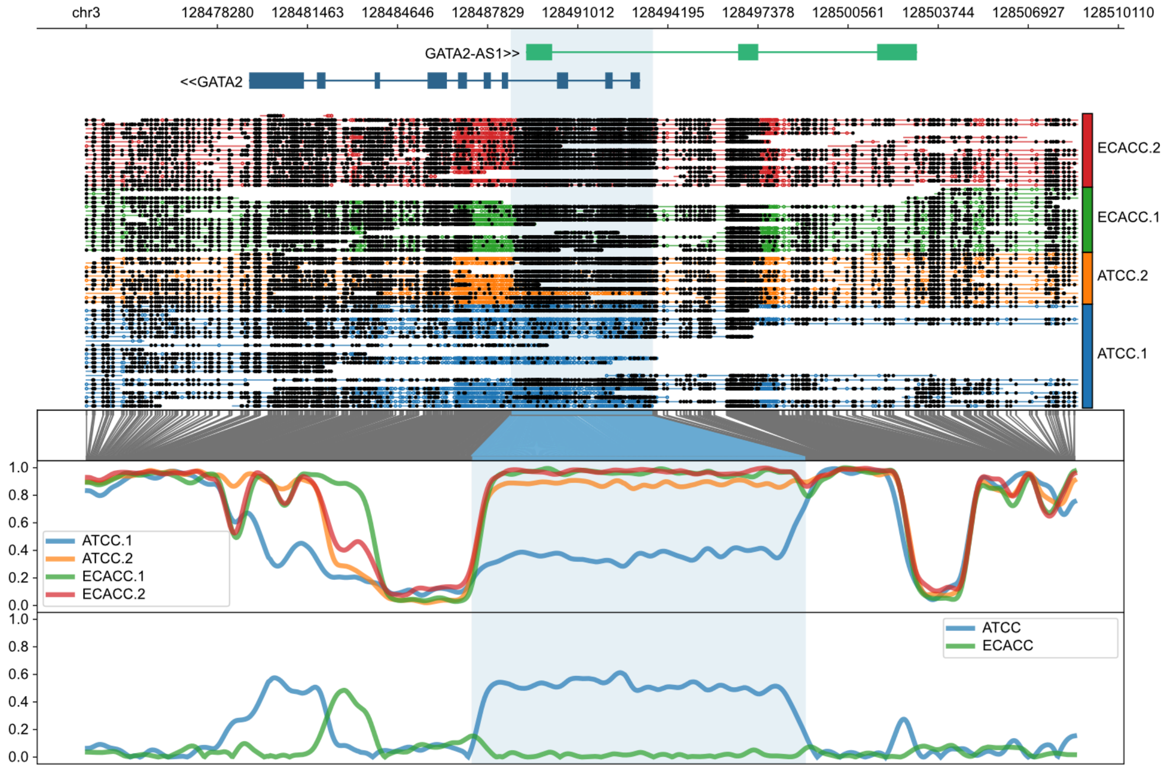


**Supplementary Figure 3b:** GATA2 allele-specific methylation pattern showing hypomethylation of one allele from the ATCC-derived sample

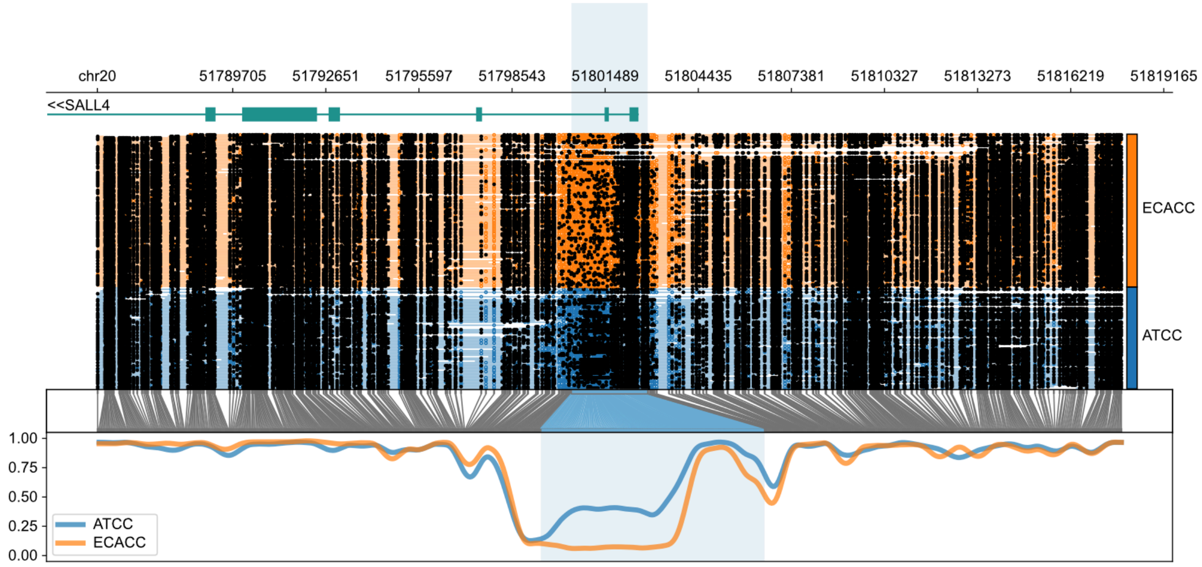


**Supplementary Figure 4a:** SALL4 differential methylation pattern, not allele-resolved

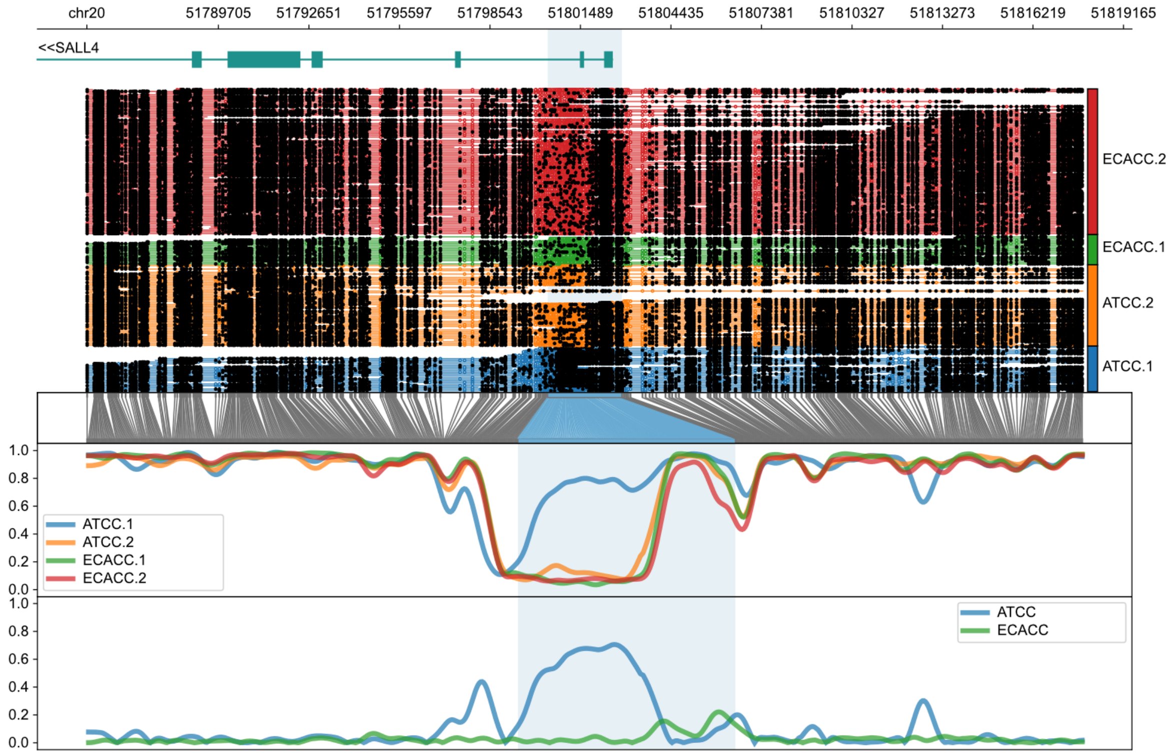


**Supplementary Figure 4b:** SALL4 allele-specific methylation pattern showing hypermethylation of one allele from the ATCC-derived sample

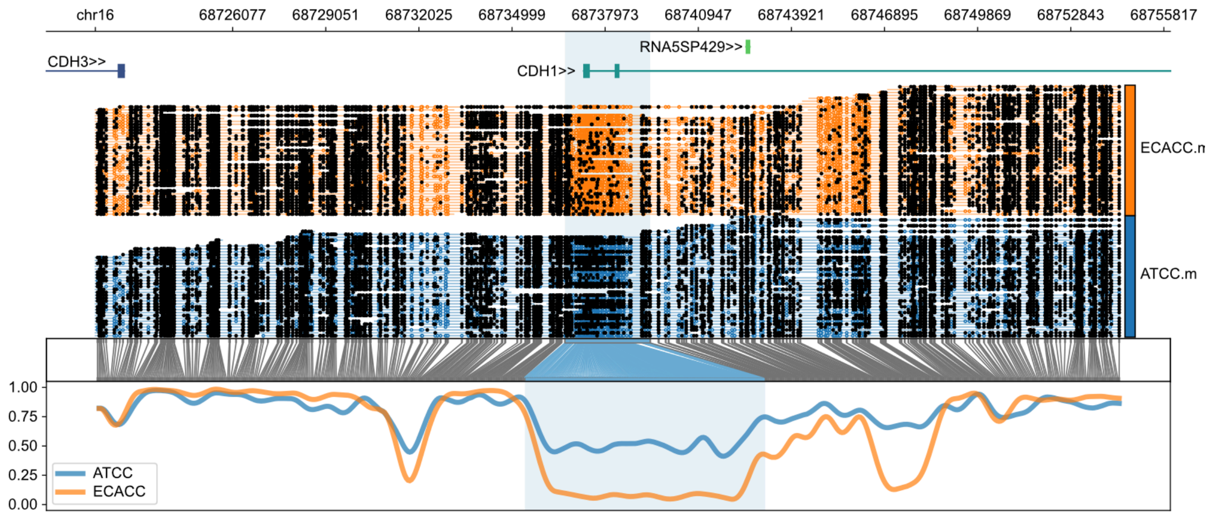


**Supplementary Figure 5a:** CDH1 differential methylation pattern, not allele-resolved

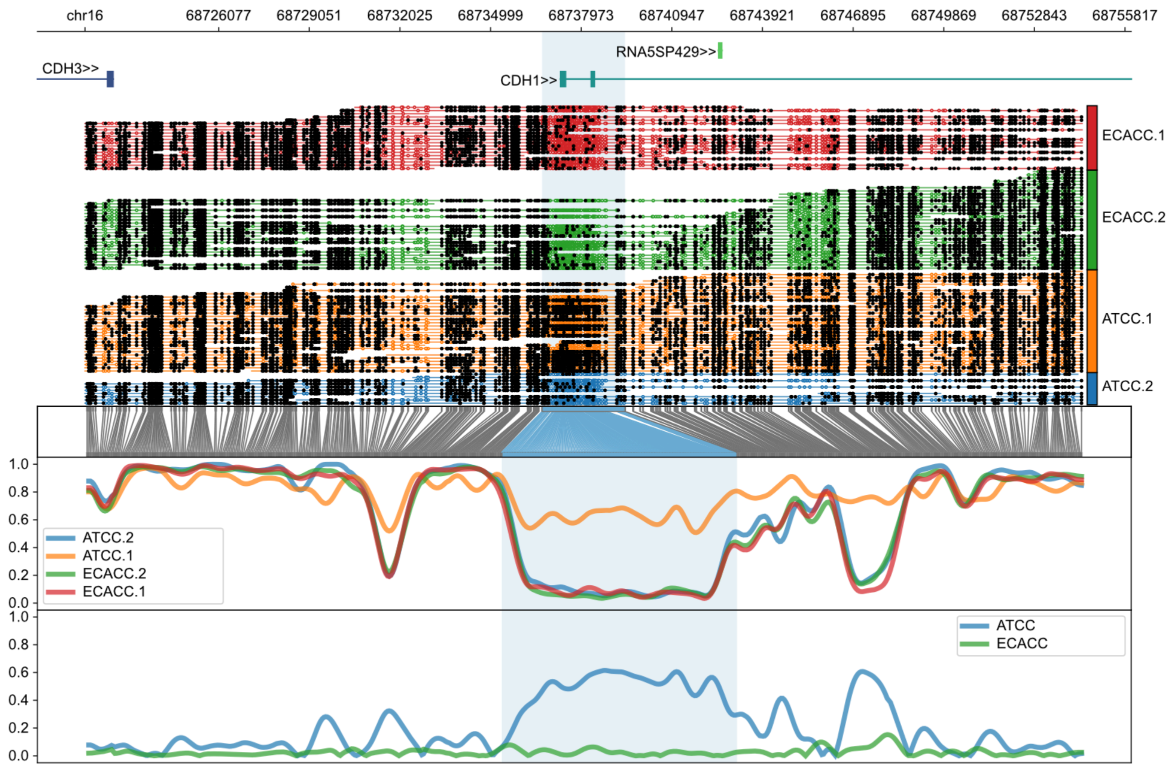


**Supplementary Figure 5b:** CDH1 allele-specific methylation pattern showing hypermethylation of one allele from the ATCC-derived sample

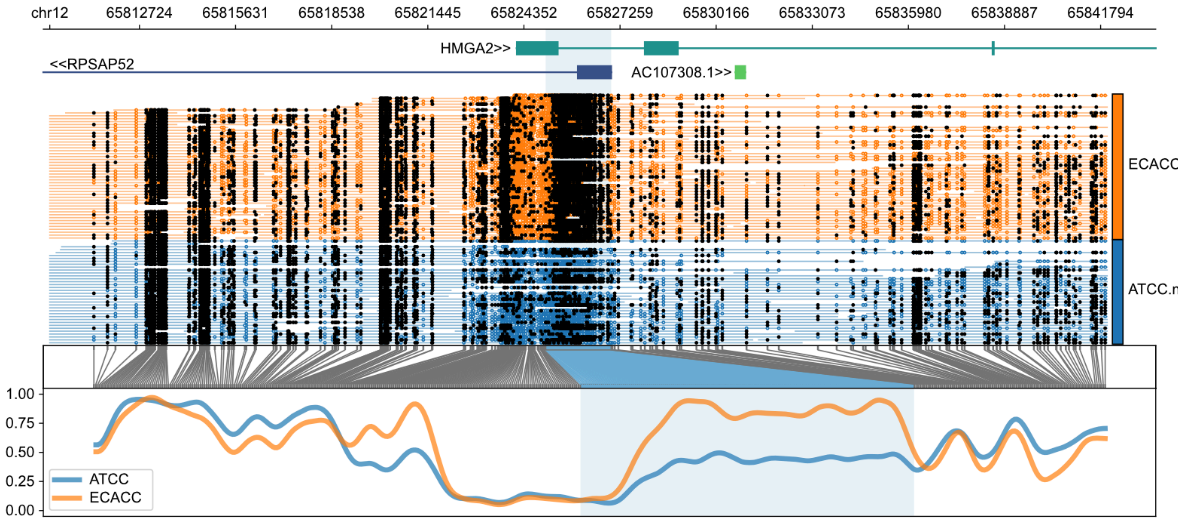


**Supplementary Figure 6a:** HMGA2 differential methylation pattern, not allele-resolved

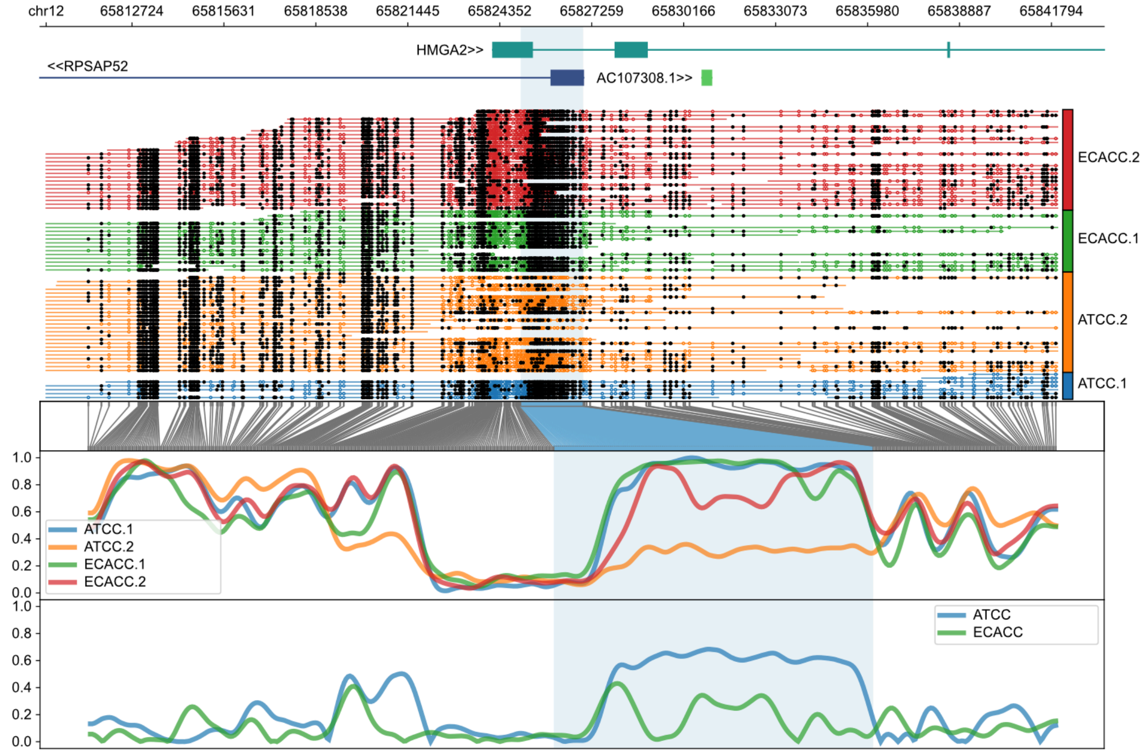


**Supplementary Figure 6b:** HMGA2 allele-specific methylation pattern showing hypomethylation of one allele from the ATCC-derived sample

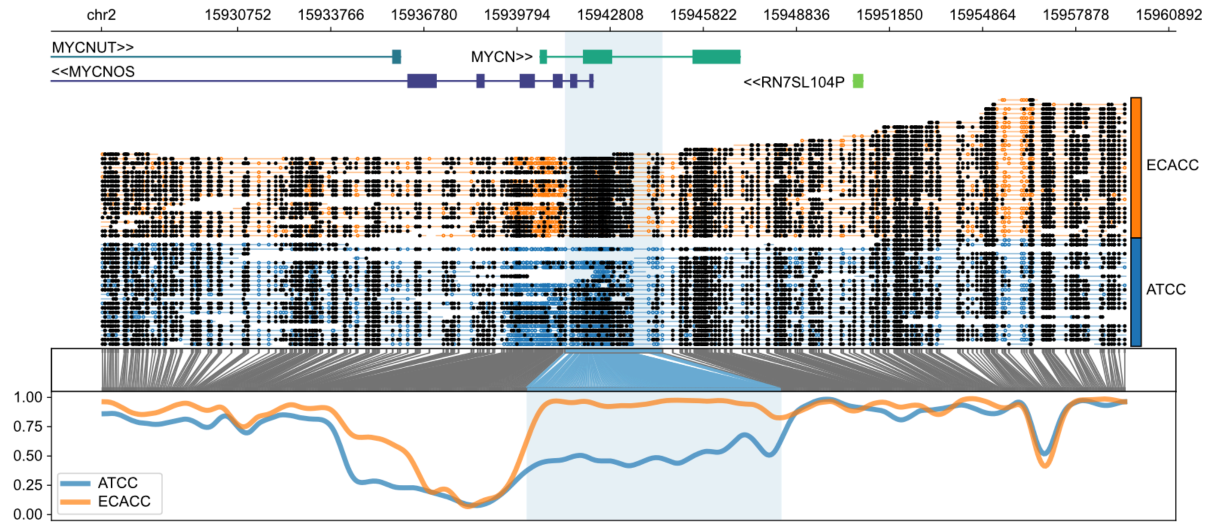


**Supplementary Figure 7a:** MYCN differential methylation pattern, not allele-resolved

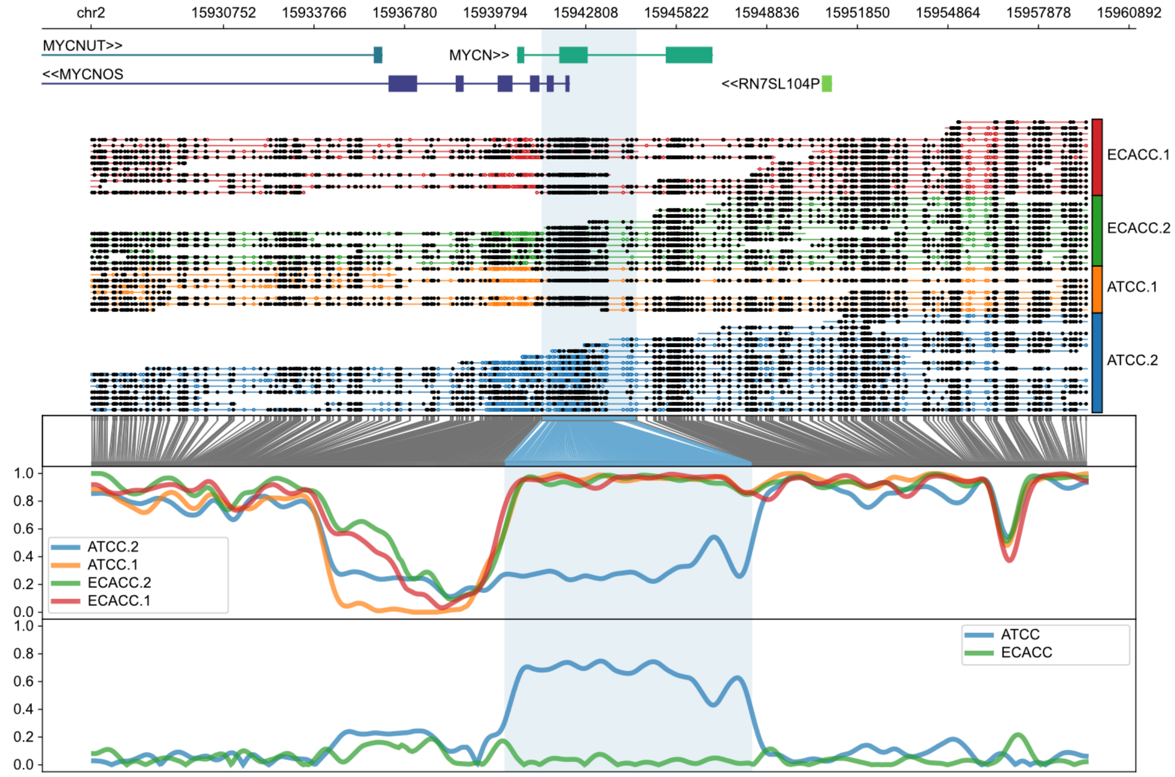


**Supplementary Figure 7b:** MYCN allele-specific methylation pattern showing hypomethylation of one allele from the ATCC-derived sample

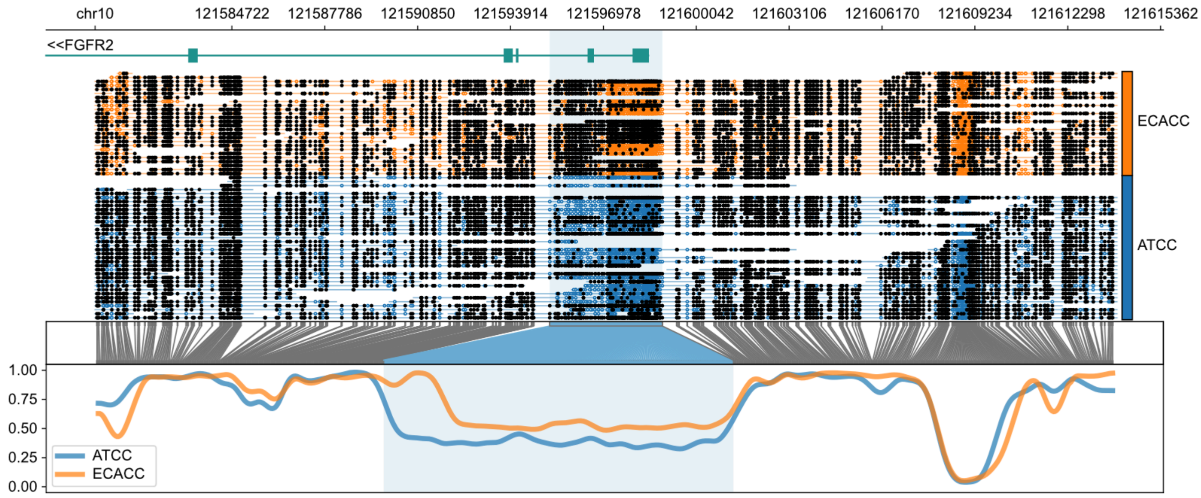


**Supplementary Figure 8a:** FGFR2 differential methylation pattern, not allele-resolved

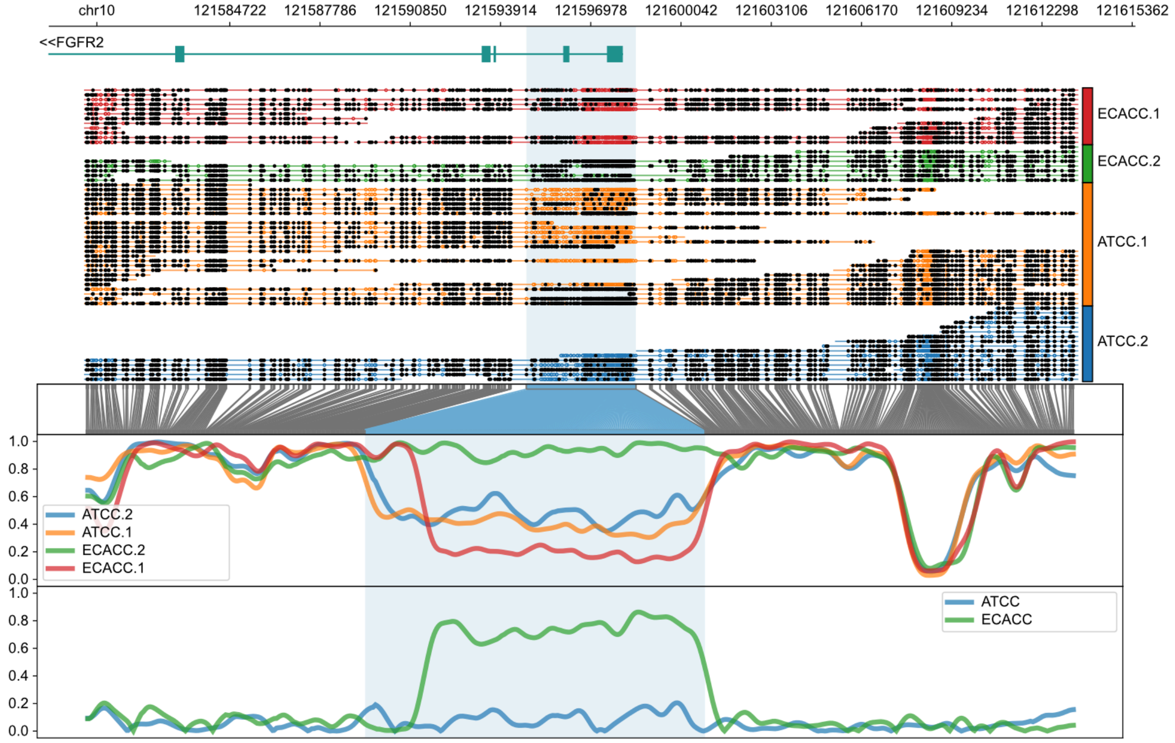


**Supplementary Figure 8b:** FGFR2 allele-specific methylation pattern showing hypermethylation of one allele from the ECACC-derived sample

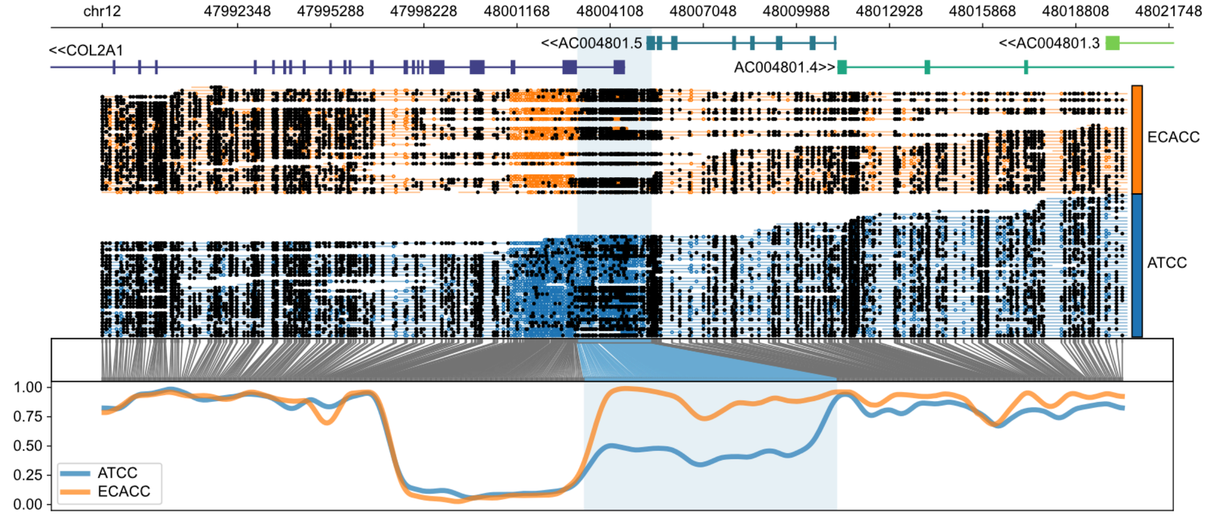


**Supplementary Figure 9a:** COL2A1 differential methylation pattern, not allele-resolved

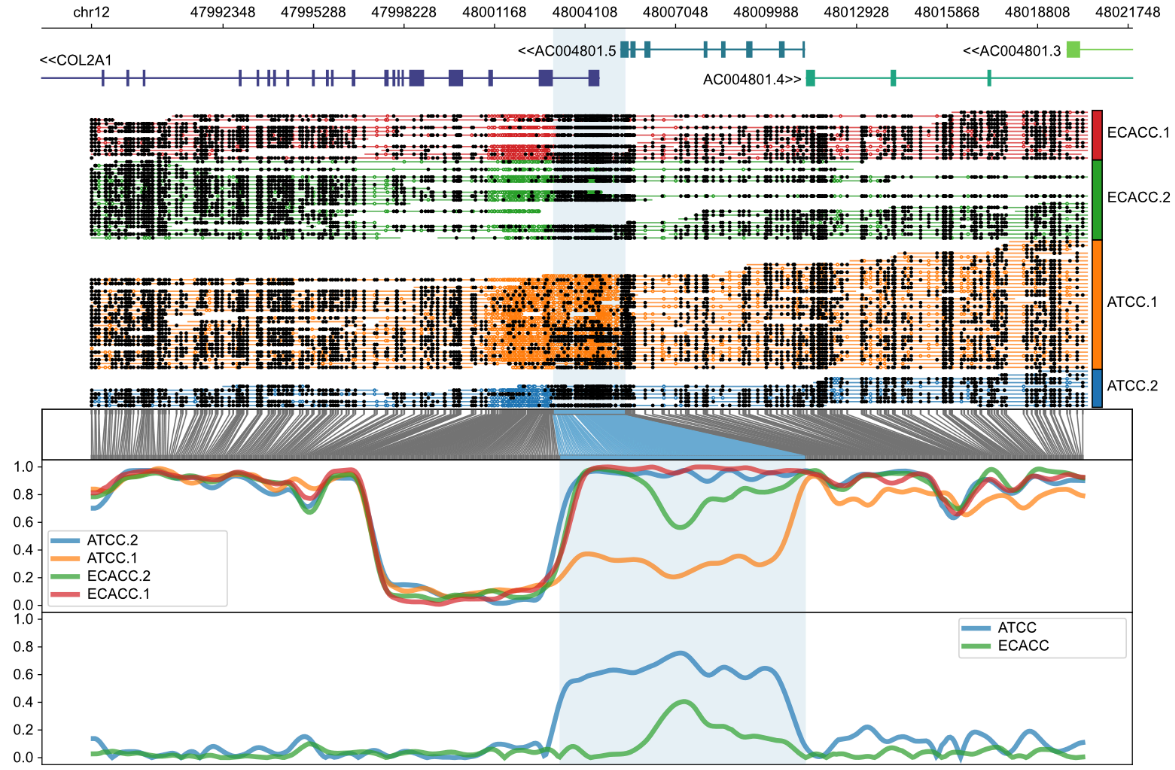


**Supplementary Figure 9b:** COL2A1 allele-specific methylation pattern showing hypomethylation of one allele from the ATCC-derived sample

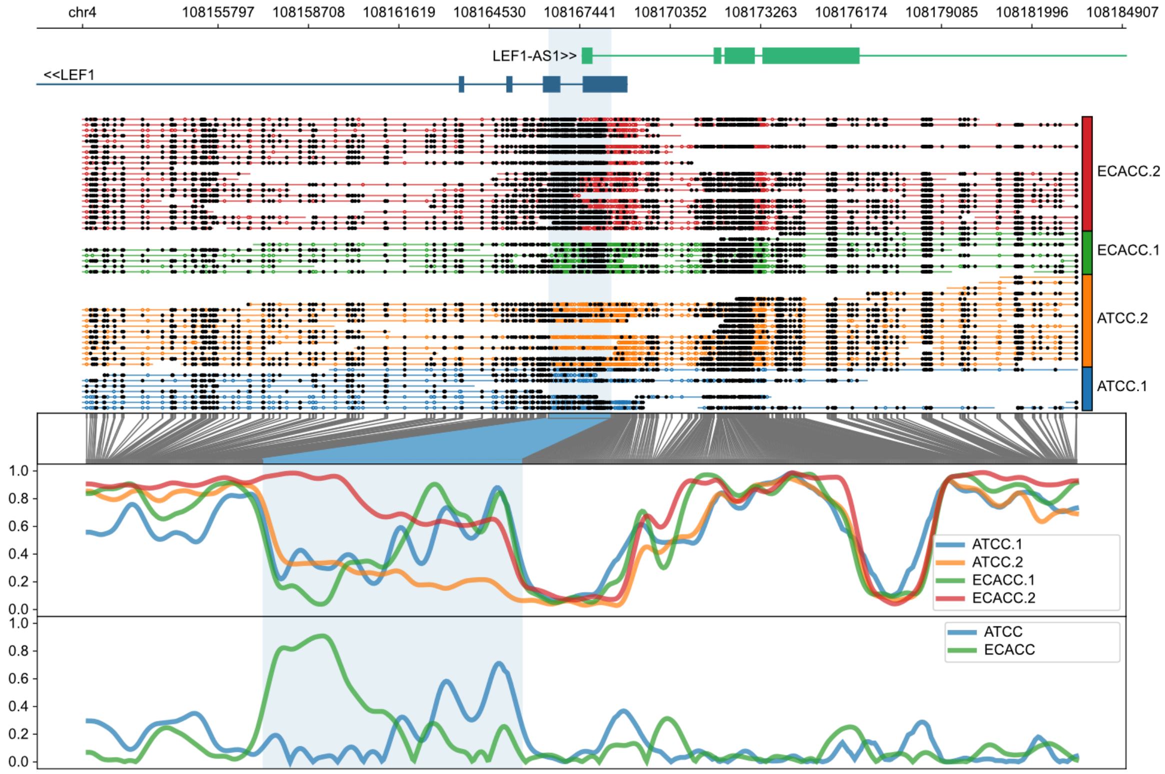


**Supplemental Figure 10:** Allele-specific differential methylation where one allele in ECACC-derived cells is hypermethylated, overlapping LEF1 and the non-coding antisense gene LEF1-AS1.

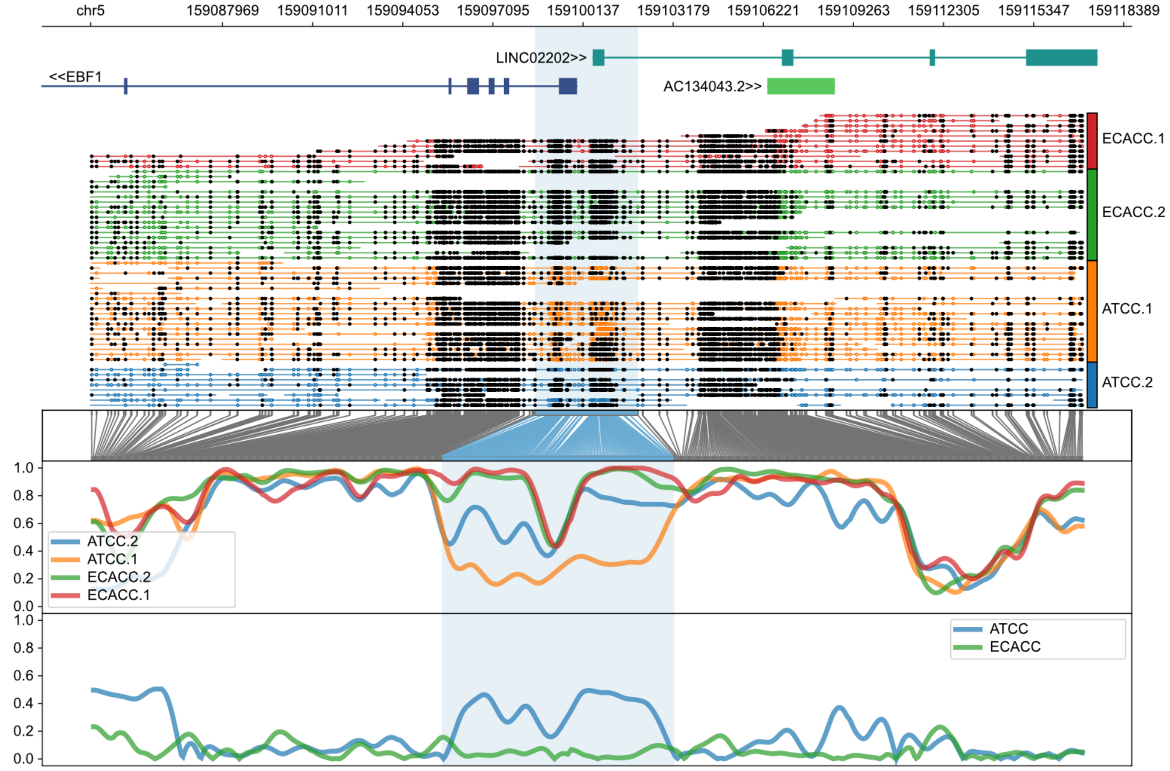


**Supplemental Figure 11:** Allele-specific differential methylation where one allele in ATCC-derived cells is hypomethylated, overlapping EBF1 and the non-coding antisense gene LINC02202.

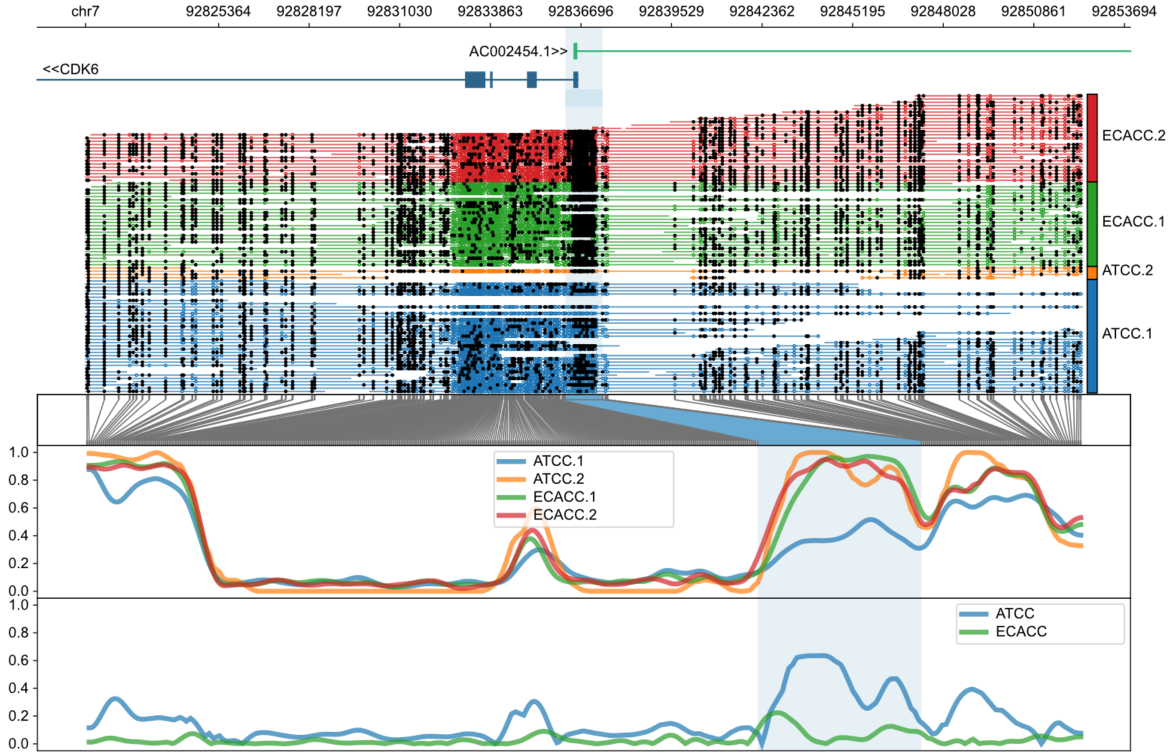


**Supplemental Figure 12:** Allele-specific differential methylation where one allele in ATCC-derived cells is hypomethylated, overlapping CDK6 and the non-coding antisense gene CDK6-AS1 (AC002454).

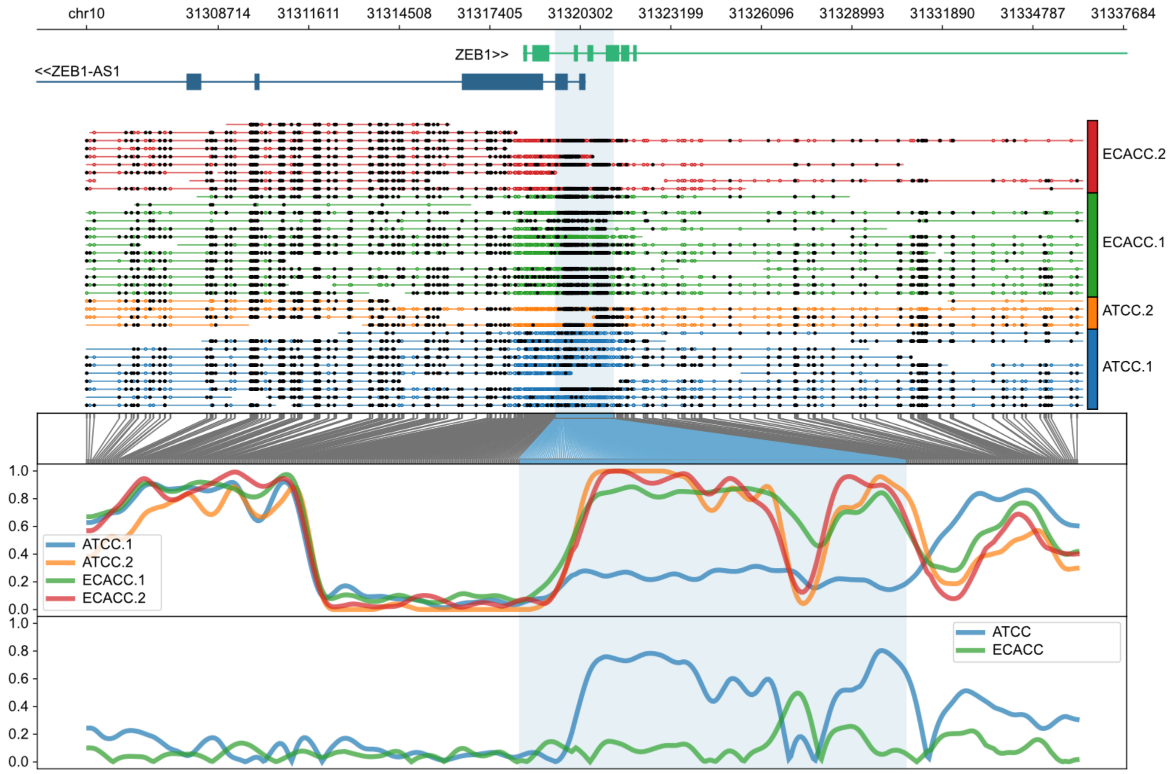


**Supplemental Figure 13:** Allele-specific differential methylation where one allele in ATCC-derived cells is hypomethylated, overlapping ZEB1 and the non-coding antisense gene ZEB1-AS1.

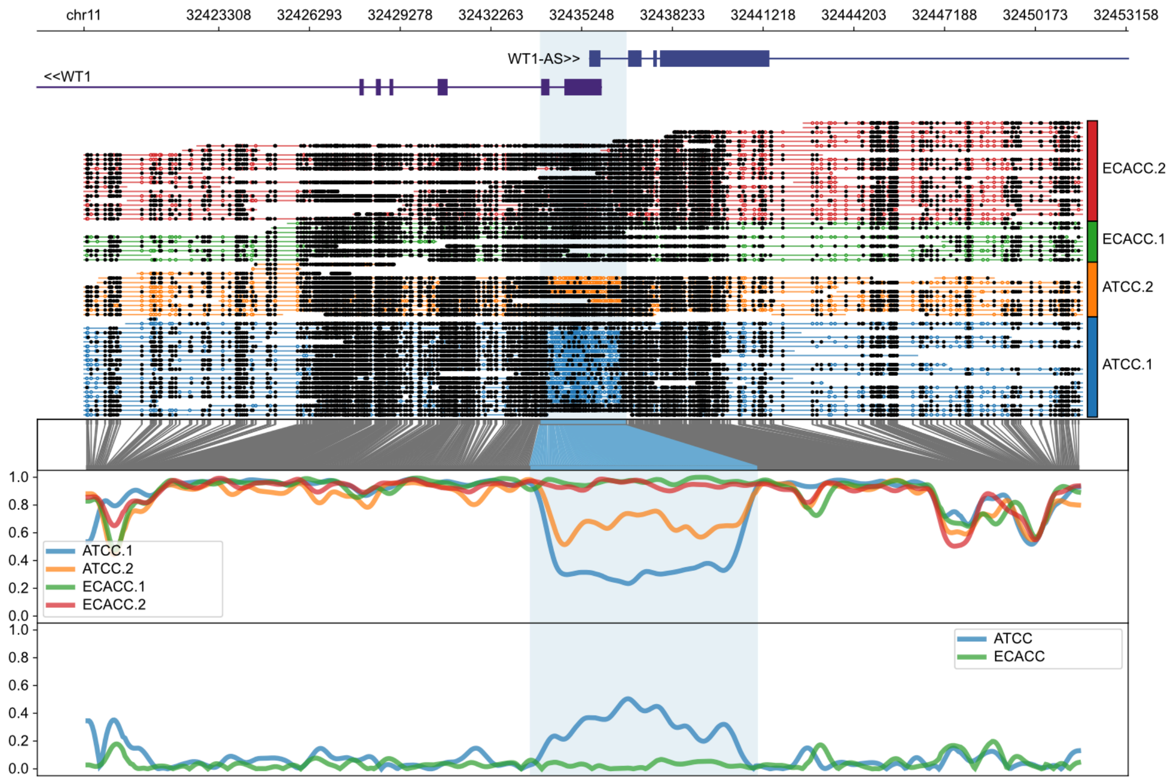


**Supplemental Figure 14:** Allele-specific differential methylation where ATCC-derived cells show inter-allele differential methylation but ECACC-derived cells do not, overlapping WT1 and the non-coding antisense gene WT1-AS.

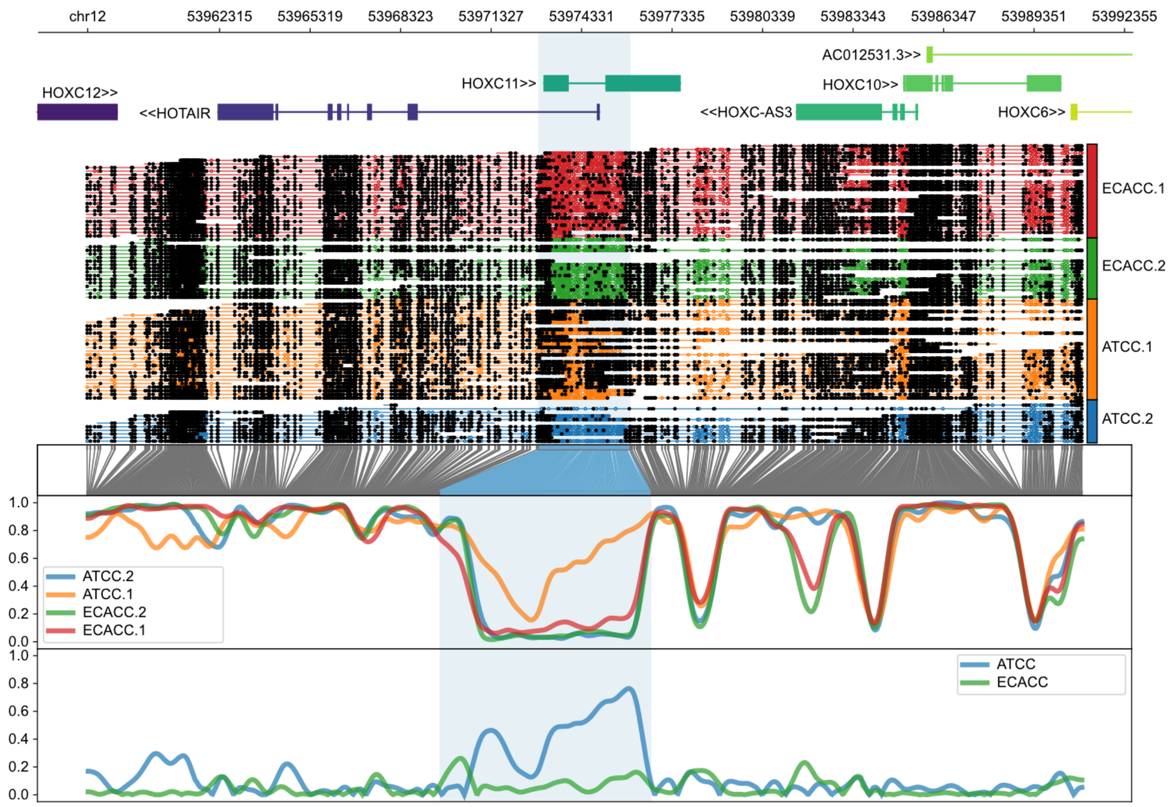


**Supplemental Figure 15:** Allele-specific differential methylation where one allele in ATCC-derived cells is hypermethylated, overlapping HOXC11 and the non-coding antisense gene HOTAIR.

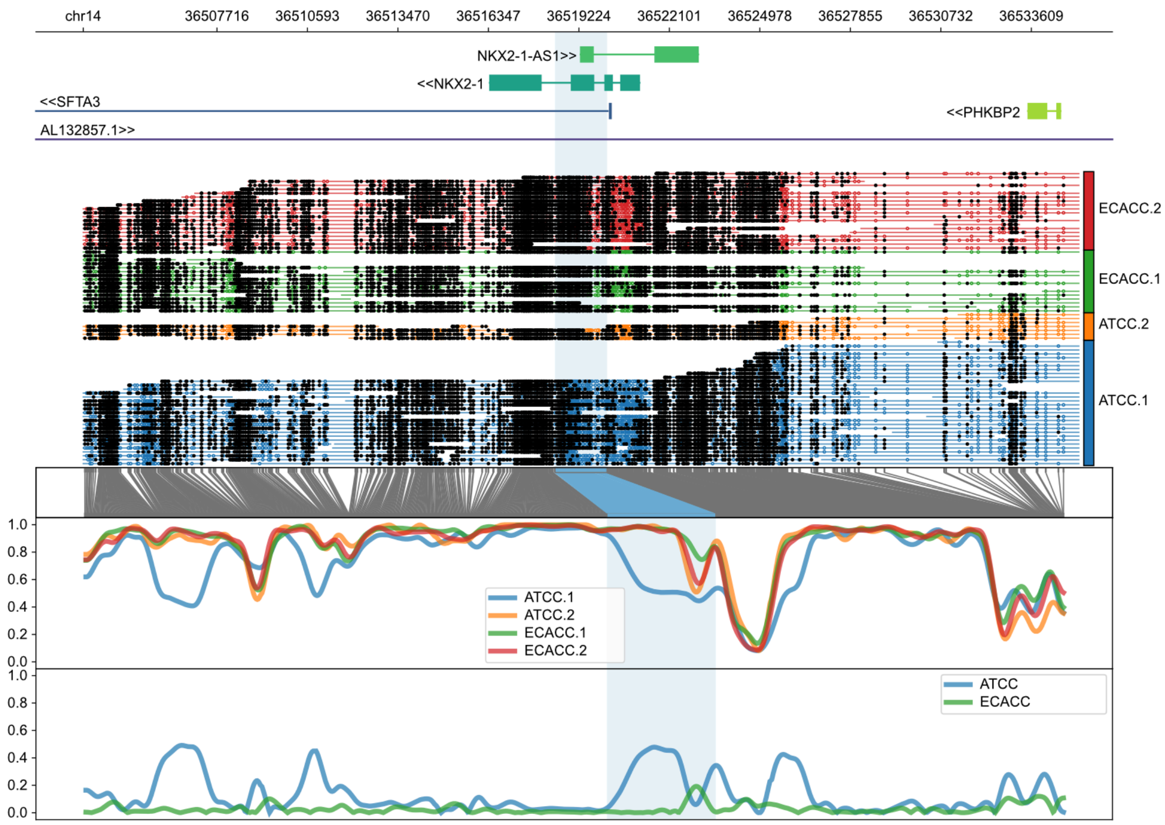


**Supplemental Figure 16:** Allele-specific differential methylation where one allele in ATCC-derived cells is hypomethylated, overlapping NKX2-1 and the non-coding antisense gene NKX2-1-AS1.

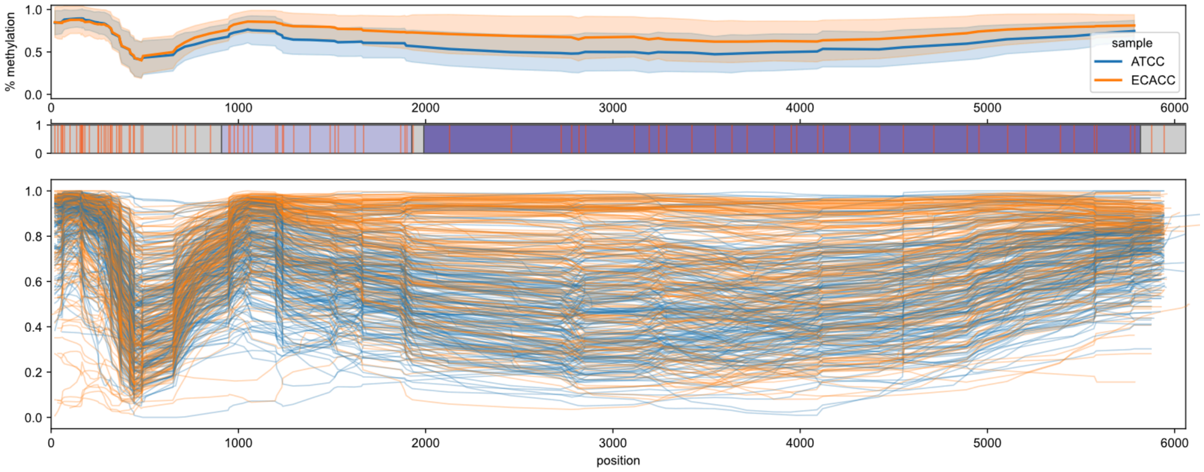


**Supplemental Figure 17:** Individual Full-length (>5900bp) L1Hs elements as annotated by RepeatMasker on hg38 via the UCSC Genome Browser. The top panel reflects the average methylation level for L1s in ATCC-derived (blue) or ECACC-derived (orange) cells +/- 95% c.i. The middle panel depicts CpG locations (red ticks) in L1 where ORF1p and ORF2p are shown as light and dark blue rectangles. The bottom panel shows the methylation profile of each individual L1 element (n=309) coloured as above.
